# The structure of pairwise competition is responsible for sudden regime shifts in a microbe-plasmid model

**DOI:** 10.1101/2025.09.08.674846

**Authors:** Ying-Jie Wang, Alberto Megías, José A. Capitán, Leonardo Aguirre, Shai Pilosof, David Alonso

## Abstract

Plasmids play a crucial role in bacterial communities by facilitating the horizontal transfer of genetic material, particularly genes that confer resistance to different stressor agents. In this work, we introduce a microbe-plasmid model to study the conditions under which a plasmid-carrying bacterial strain can invade its corresponding plasmid-free counterpart, as well as those under which it can be eradicated from the population through potential intervention strategies. We reveal the variety of dynamic regimes of the model depending on the parameter values. In particular, we show that a heterogeneous structure of the competition between the two strains drives the possibility of coexistence of stationary states (i.e. bistability) which creates conditions for sudden regime shifts and hysteresis. We explore the implications of our results for the potential eradication of plasmid-mediated antibiotic resistance.

## 1 Introduction

Plasmids play a crucial role in bacterial communities by facilitating the horizontal transfer of genetic material, particularly genes that confer resistance to different *environmental* stressors, such as antibiotics and heavy metals [1–3]. The global burden of antibiotic resistance (ABR), largely driven by plasmid-mediated gene transfer, poses a serious threat to public health and undermines the effectiveness of current treatment strategies [4, 5]. These extrachromosomal DNA molecules enable bacteria to rapidly adapt to selective pressures, such as antibiotic exposure, by sharing resistance traits across diverse species and environments. This ability accelerates the spread of multidrug-resistant pathogens, which poses a significant challenge to public health and antimicrobial treatment strategies [6–8]. Therefore, plasmids contribute to the persistence and evolution of resistance genes in microbial communities, underscoring their importance in the ongoing battle against antibiotic resistance.

Models of microbe-plasmid dynamics have traditionally adopted a two-dimensional (2D) framework that tracks the population dynamics of plasmid-free and plasmid-carrying strains (subpopulations). This modeling approach has been widely applied to investigate plasmid coinfection [9], the removal of target plasmids via denial-of-spread (DoS) strategies [10], mechanisms of plasmid stabilization in polyclonal microbial communities [11], the selection for adaptive immune systems [12], and the role of horizontal gene transfer in promoting microbial community stability [13]. Despite these important advances, previous studies have not investigated the full range of dynamic regimes (alternative states [14])—the stationary states in which one strain, both strains, or neither strain persists—nor the regime shifts that occur across parameter space.

Moreover, current models typically neglect direct intraspecific competition, i.e., the interference competition [15, 16] within a population. Such competition can result in cell death, for instance, driven by plasmid-encoded toxin-antitoxin (TA) systems that selectively eliminate plasmid-free cells, or through TA-mediated self-regulation under stress or resource limitation [17–19]. However, the impact of direct intraspecific competition on key ecological outcomes—such as the ability to prevent plasmid invasion into plasmid-free populations (plasmid invasibility) or to eliminate plasmids from plasmid-established populations (plasmid eradicability)—remains poorly understood. Bridging these knowledge gaps may offer new insights into context-dependent strategies for controlling the spread of plasmid-encoded antibiotic resistance.

In this work, we introduce a 2D microbe-plasmid model to investigate the conditions under which a plasmid-carrying strain can invade a plasmid-free population, as well as the conditions under which a plasmid can be eradicated from the population. We characterize the different model dynamic regimes depending on model parameter values. When two individual bacterial cells encounter each other, they can either initiate a conjugation process or one cell can kill the other. The latter interaction represents an encounter-induced death of one of the two cells, which decreases the overall competition pressure in the population. We label this type of process “direct competition” or “pairwise competition-induced death”. First, we study how the variety of dynamic regimes depends on model parameters and, crucially, on the type of competition and the structure of the competition matrix. Then, we derive sufficient conditions for the invasion of the plasmid-carrying bacterial strain. Finally, we compare different eradication strategies in the presence (or not) of a bistationary state (i.e., the possibility of coexistence of two stationary stable states [20]). We describe conditions under which sudden regime shifts and hysteresis can arise as control parameters vary. These findings have broad implications for understanding plasmid introduction and persistence in bacterial communities, and are particularly relevant for designing effective strategies to eradicate antibiotic resistance.

## 2 The Model

The model considers a plasmid-free bacterial population growing in a resource-limited environment through the processes of cell division and death. In addition, bacterial cells, which all belong to the same bacterial population, can carry a plasmid, which we define as the plasmid-carrying strain. The one-bacterial population-one-plasmid (1B1P) model considers then the competition dynamics between the two strains—the plasmid-bearing and plasmid-free strains— when growing together. The dynamics of the two strains are described by the following five elemental processes (see Table 1):

**Table 1:** The dynamics of the plasmid-free strain (left column) and the plasmid-carrying strain (right column). A plasmid-free cell, *A*_0_, and a plasmid-carrying cell, *A*_1_, require a common resource for growth, represented here by the empty space. Cell division occurs at different rates (*β*_0_ *> β*_1_). Pairwise cell encounters take place at rate *γ*_0_. Death for both strains occurs at different rates (which are only the same if the resistance parameter *r* is 0). After an encounter takes place, there is a chance that conjugation will occur (probability *χ*), or, alternatively, one of the two cells may die due to competition (probabilities *p*_00_, *p*_01_, *p*_10_, *p*_11_), thereby creating available space (resources). Finally, plasmid-carrying cells may fail to correctly segregate the plasmid into the two daughter cells (with probability *ϵ*).

| Process | Reaction | Rate | Reaction | Rate |
| --- | --- | --- | --- | --- |
| **Cell Division** | $A_0 + \emptyset \rightarrow A_0 + A_0$ | $(\beta_0)$ | $A_1 + \emptyset \rightarrow A_1 + A_1$ | $(\beta_1(1 - \epsilon))$ |
| **Cell Death** | $A_0 \rightarrow \emptyset$ | $(\delta_0 + \delta_1)$ | $A_1 \rightarrow \emptyset$ | $(\delta_0 + \delta_1(1 - r))$ |
| **Competition** | $A_0 + A_1 \rightarrow A_1 + \emptyset$ | $(\gamma_0 p_{01})$ | $A_0 + A_1 \rightarrow A_0 + \emptyset$ | $(\gamma_0 p_{10})$ |
| | $A_0 + A_0 \rightarrow A_0 + \emptyset$ | $(\gamma_0 p_{00})$ | $A_1 + A_1 \rightarrow A_1 + \emptyset$ | $(\gamma_0 p_{11})$ |
| **Segregation error** | | | $A_1 + \emptyset \rightarrow A_1 + A_0$ | $(\beta_1 \epsilon)$ |
| **Conjugation** | $A_1 + A_0 \rightarrow A_1 + A_1$ | $(\gamma_0 \chi)$ | $A_0 + A_1 \rightarrow A_1 + A_1$ | $(\gamma_0 \chi)$ |

1. **Cell Division**. Each bacterial cell divides into two daughter cells with rates *β*_0_ and *β*_1_. Generally, cells without plasmids divide more quickly (*β*_1_ *< β*_0_), as plasmid carriage is considered a cost that slows cell division. We express *β*_1_ = *β*_0_(1 *− c*), where *c* denotes the *cost* and ranges between 0 and 1. Errors in plasmid segregation during cell division can occur with a probability *ϵ*. Empirical studies suggest that the probability of such segregation errors is low [21]. We also introduce the logistic behavior of cell division, representing competition on a shared, limited resource that slows down the division rate as the community size increases towards the carrying capacity [21, 22].
2. **Cell Death**. This type of mortality is independent of population density, affecting every cell equally at a constant per capita rate. Plasmid-carrying cells exhibit reduced mortality rates when certain selective pressures are applied. This phenomenon is represented by the parameter *r*, which ranges between 0 and 1. In the presence of stress factors, mortality rates escalate by *δ*_1_. Nevertheless, if *r* achieves its maximum value of 1, plasmid-carrying cells remain unaffected by stress. The rate *δ*_0_ denotes the baseline per capita mortality rate in an environment devoid of stressors.
3. **Competition-induced Death**. Every cell can negatively impact other individual cells, such as via allopathic compounds. This impact might occur between cells of the same strain (self-inhibition) or cells of different strains (cross-inhibition). However, it is generally stronger on cells of the same strain. Here, we study the general case, where carrying a plasmid can affect (negatively or positiverly) both cross- and self-inhibition. These types of intra- and inter-strain inhibitions are modeled by mortality induced by encounters between pairs (at a rate *γ*_0_). Once an encounter has taken place between two cells, there is a probability *p*_*ij*_ of a death to occur in the *i*-th cell (*i* = 0, 1) caused by the *j*-th cell (*j* = 0, 1). The four *p*_*ij*_ elements (*p*_00_, *p*_01_, *p*_10_, *p*_11_) define the structure of the direct competition matrix.
4. **Segregation Error** When plasmid-carrying cells divide, there is a chance that the plasmids might not be distributed between the two resulting daughter cells. As a result, one daughter cell might end up without any plasmids, known as a plasmid-free cell. This occurrence has a low probability, denoted by *ϵ*.
5. **Conjugation**. Each plasmid-carrying cell is prone to infect other cells that do not contain this plasmid. This process requires first the encounter of the two cells, at rate *γ*_0_. Once this has happened, there is a probability *χ* of a conjugation to occur.

The four dynamic rates define unique time scales: the pace of cell growth (*β*_0_, growth timescale), the rate at which cells move and interact (*γ*_0_, encounter timescale), the lifespan of cells in regular conditions (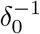, natural death timescale), and in the presence of environmental stressors ((*δ*_0_ + *δ*_1_)^*−*1^, stress-induced death timescale). Naturally, these time scales may overlap. The rest eight parameters (a conjugation probability, *χ*, 2*×*2 direct competition matrix, plasmid cost and resistance parameters, *c* and *r*, and a segregation error *ϵ*) range between 0 and 1.

## 3 Results

### Model Analysis

The five processes mentioned above create a competitive dynamic between the plasmid-free and plasmid-carrying strains due to limited resources, which means that there is a limit on the number of cells that can occupy a given volume or surface area. Consequently, cells compete for the available *free space*-associated resources. This upper limit on cell density (i.e. carrying capacity) is designated by the parameter *K*. When *K* is low, so does individual abundance, and therefore stochastic factors significantly influence outcomes. In this study, we focus on the scenarios where *K* is high, allowing us to analyze the dynamics using the following 2D ODE system:

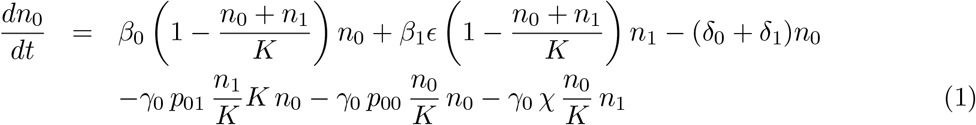

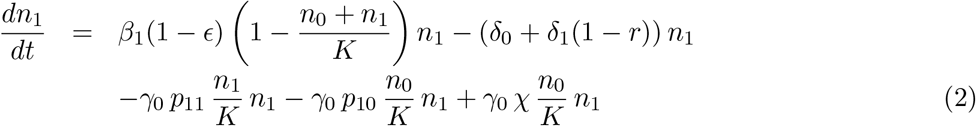

For convenience, we divide the two equations by *K* and *γ*_0_, which creates an equivalent 2D system, where a dimensionless time *τ* should be defined:

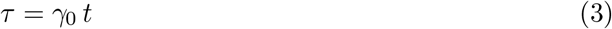

This causes the remaining rates to be given as dimensionless rates (or ratios) with respect to *γ*_0_ (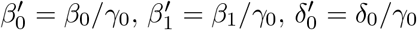, and 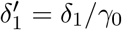). The system reads:

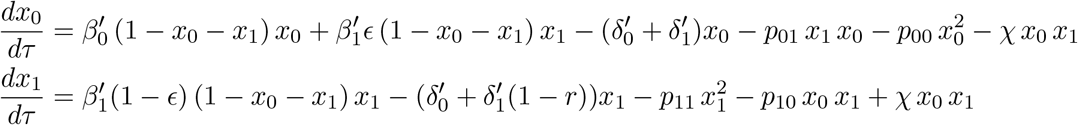

Moreover, two novel densities emerge relative to the system carrying capacity *K*, characterized by *x*_0_ = *n*_0_*/K* and *x*_1_ = *n*_1_*/K*. Henceforth, we omit the apostrophe from the rates and designate 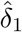 as *δ*_1_ (1 *− r*), resulting in:

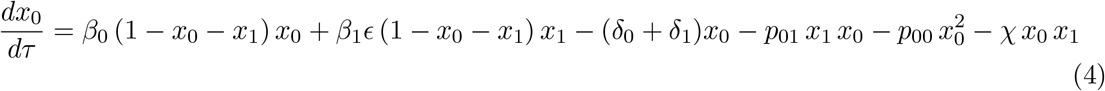

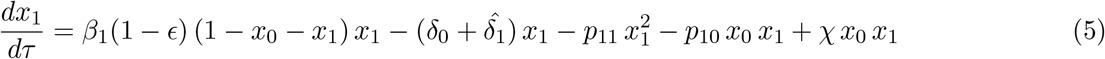

The system becomes more compact by typically defining the two following functions:

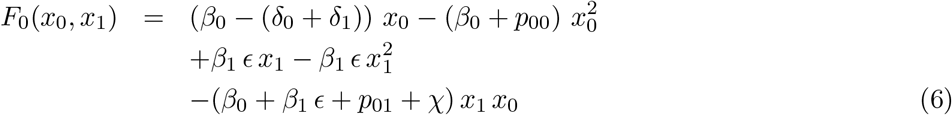

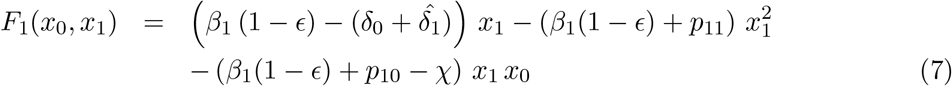

which results in:

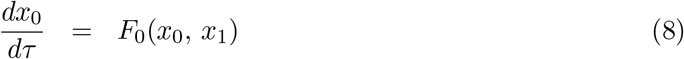

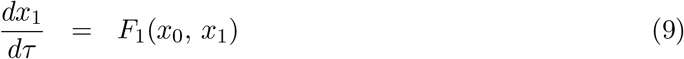

The Jacobian matrix of the system is then defined as

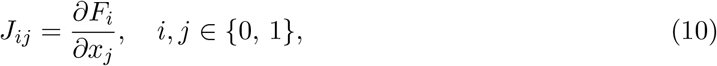

and can also be calculated, which will be of much use in further analyses:

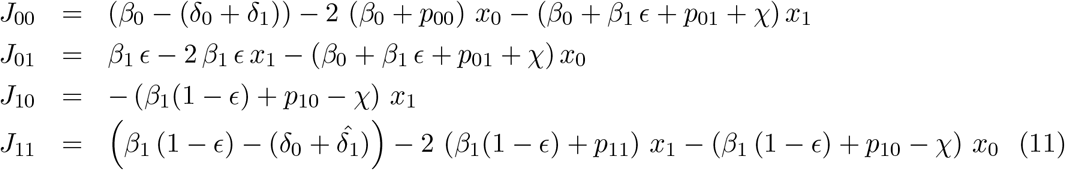

### Stationary Points

In a standard manner, these points can be calculated by solving for *x*_0_ and *x*_1_ in the following 2D system of non-linear equations:

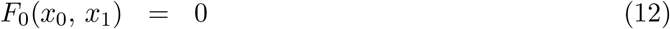

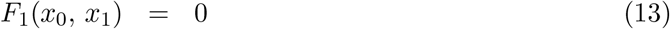

Simple inspection of this system reveals that the resting points of the dynamics can be of three types:

1. System-empty equilibrium. If *x*_0_ and *x*_1_ are both zero, the pair (0, 0) is a trivial solution of Eqs (12)-(13).
2. Plasmid-free equilibrium. In the absence of plasmid-carrying cells (*x*_1_ = 0), the system exhibits a logistic behavior toward this equilibrium:

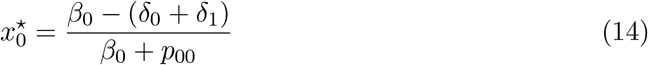
3. Coexistence equilibrium. This equilibrium is characterized by non-zero densities of both strains. By assuming it exists, i.e., 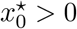 and 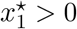, then we can take the 2nd equation of the system (Eq 13):

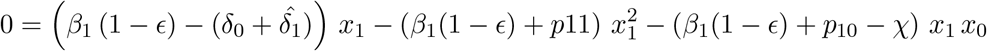

and divide it by *x*_1_:

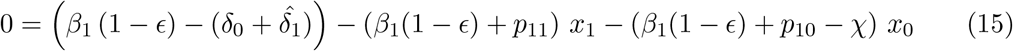

The solution of the system of two equations—Eqs. (12) and (15)—gives rise to a quadratic equation. In order to express it in a simple way, we introduce the following intermediate parameters:

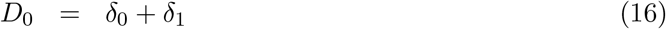

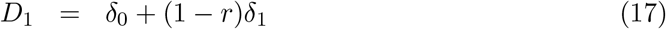

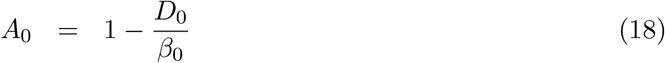

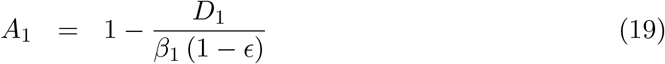

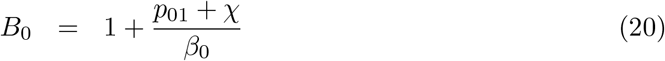

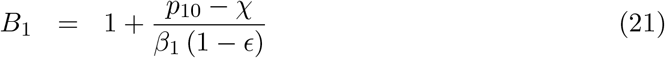

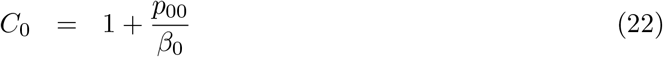

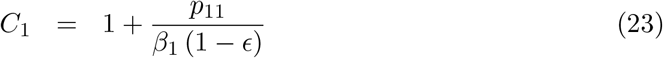

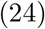

Now, we can use Eq (15) to express *x*_0_ in terms of *x*_1_:

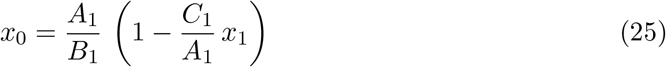

If we introduce *x*_0_ (in terms of *x*_1_) into Eq (12) and express the resulting equation in terms of an intermediate variable *η*, defined as *C*_1_*x*_1_*/A*_1_, we obtain a simple quadratic equation:

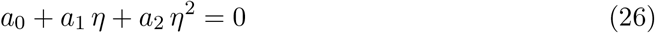

where the three coefficients are:

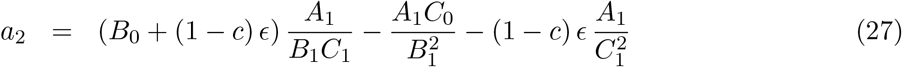

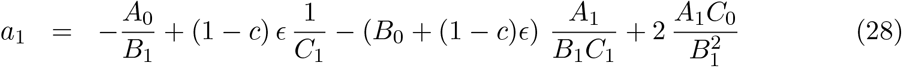

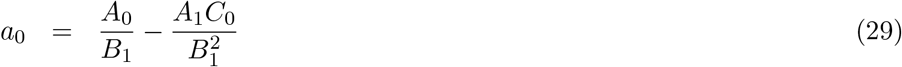

Given that

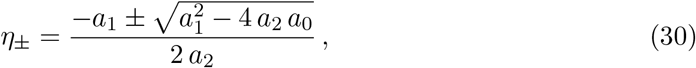

the two solutions for *x*_1_ and *x*_0_ can be written as:

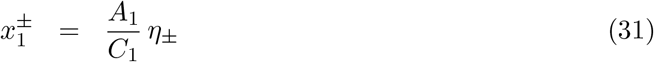

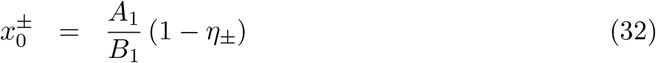

### Parameter limitation

Notice that, by definition, there are some constraints on the parameters we have used. In particular, if *γ*_0_ is the rate of encounter of a pair of individual cells, then, upon encounter, three outputs are possible:

1. The death of one particular cell of the pair complex, with probabilities *p*_00_ and *p*_11_ for conspecific encounters, and *p*_01_ and *p*_10_ for heterospecific encounters.
2. In heterospecific encounters, conjugation can take place, with probability *χ*.
3. Nothing happens, with probability (1 *− p*_00_) or (1 *− p*_11_) for conspecific encounters of plasmid-free and plasmid-carrying cells, respectively, and with probability (1*−p*_01_*−p*_10_*− X*) for heterospecific encounters. Therefore, since all these are probabilities, *p*_00_, *p*_01_, *p*_10_, *p*_11_ and *χ* have to meet the following contraints: *p*_00_ *<* 1, *p*_11_ *<* 1, and *p*_01_ + *p*_10_ + *X* < 1.

### Plasmid Invasibility

Since plasmids frequently contain genes for antibiotic resistance along with other functions, it is worth investigating the circumstances that allow a plasmid-carrying strain to penetrate a plasmid-free population that has stabilized in isolation.

This condition can be assessed by looking at the dynamics of small perturbations of *x*_1_ when the *x*_0_ rests on its equilibrium point. In this situation, as *x*_1_ ≪ 1, its early dynamics is given by:

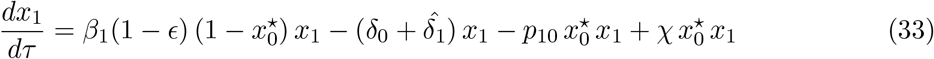

which can be written as:

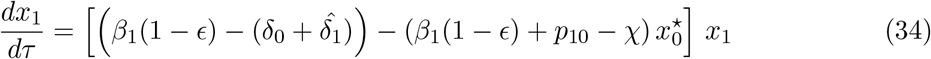

Plamid invasion will be possible if:

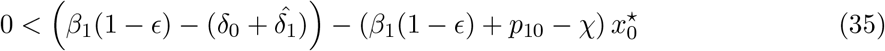

This condition can be rewritten as:

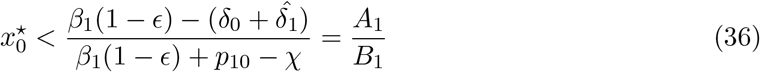

This equation sheds light on the conditions that support plasmid invasion. In particular, it allows us to establish two sufficient criteria corresponding to two possible scenarios: either both the numerator and the denominator in Eq. (36) are positive, or both are negative, thereby maintaining the positive value of the ratio *A*_1_*/B*_1_. Then, in these cases, two simple sufficient conditions for plasmid invasion can be respectively derived (since both involve that the right-hand side of the inequality in (36) is larger than 1 (*A*_1_ *> B*_1_):

1. Both numerator and denominator are positive. If *β*_1_(1 *− ϵ*) *> χ − p*_10_, then it is sufficient to require:

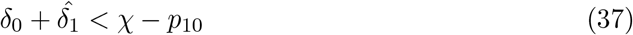
2. Both numerator and denominator and negative. If *β*_1_(1 *− ϵ*) *< χ − p*_10_, then it is sufficient to require:

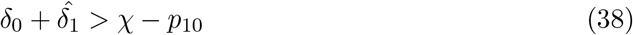

When *p*_10_ = 0 (i.e., the competition impact of plasmid-free strain *x*_0_ on plasmid-carrying strain *x*_1_ is absent), the plasmid invasion threshold is further simplified: either 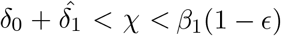, or 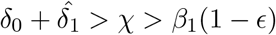. Remember that these conditions are only sufficient. Less strict conditions might exist where plasmid invasion could still occur. These sufficient conditions are noteworthy because they guarantee, in any situation, that the general condition outlined in Eq (36) is satisfied irrespective of the initial equilibrium level of the plasmid-free strain, 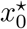. According to these conditions, increasing *p*_10_ will reduce the ranges of *χ* that allow for plasmid invasion.

### Dynamic regimes across competition scenarios

In the study of bacterial communities, understanding how different dynamic regimes manifest is crucial to predicting the fate of coexisting strains. This section delves into the dynamic regimes observed in a model that assesses the growth of a plasmid-free strain alongside its a plasmid-carrying strain. By varying parameter values, the model reveals different potential outcomes: the collapse and extinction of both strains, the persistence of only the plasmid-free strain, or the stable coexistence of both strains. Characterizing these regimes not only provides insight into the complex interactions that influence bacterial survival and adaptation, but also allows us to reveal critical thresholds in parameter values that mark a qualitative change in system behavior. In this framework, these thresholds are referred to as critical transitions, sudden changes, or regime shifts in ecological research [14], or more plainly as phase transitions in the field of statistical physics.

In general, the different dynamic regimes may not depend on all parameter values because some may only influence transient behavior. This is the case of *γ*_0_. In the dimensionless version of the model, *γ*_0_ plays no role but rescaling the time. The different regimes reported here correspond to the long-term behavior of the system. In fact, the question is, after discarding the transient, what is the qualitative state of the system once it has reached a steady state and how does it depend on model parameters? This question can be answered by (1) investigating the probability of the system reaching different dynamic regimes (i.e., extinction, E, plasmid-free state, PFS, coexistence, C, and bistationary state, BS, where a mixture of stable coexistence and stable plasmid-free state exists depending on the initial state) with a given set of parameter space, and (2) carefully analyzing the type of stability of the different stationary states.

After introducing the dimension-less model in Eqs (4)-(5), we know that these regimes should depend only on three independent dimensionless rates (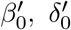, and 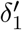), and 8 dimensionless parameters (*ϵ, c, r, p*_00_, *p*_01_, *p*_10_, *p*_11_, and *χ*, from which *ϵ*, the segregation error, is expected to be very small) that can potentially range from 0 to 1 (where *p*_10_, *p*_11_ and *χ* should, in addition, fulfill the constraints given by *p*_01_ + *p*_10_ + *X <* 1). We therefore define the parameter space of these rates and parameters for iterations (Table 2).

**Table 2:** Model parameter space for its exploration through the iterative process that generates a number of random parametric configurations withing these boundaries. Note that *β*_1_ derives from *β*_0_ (*β*_1_ = *β*_0_(1 − *c*)), and that *K* and *γ*_0_ are not considered for they make no influence on the different dynamic regimes highlighted here. All parameters reported here are the ones before being scaled by *γ*_0_ ∈ (0, 10).

|  |  |  |
| --- | --- | --- |
| Strain-specific death rate | $\delta_0$ | (0, 10.0) |
| Stress-induced death rate | $\delta_1$ | (0, 10.0) |
| Cell division rate | $\beta_0$ | (0, 10.0) |
| Segregation error probability | $\epsilon$ | (0, 0.01) |
| Plasmid transmission probability | $\chi$ | (0, 1.0) |
| Competition induced death probability | $p_{ij}$, $i, j \in \{0, 1\}$ | (0, 1.0) |
| Plasmid-specific reproduction cost | $c$ | (0, 1.0) |
| Plasmid-specific resistance | $r$ | (0, 1.0) |

For the setting of competition parameters (*p*_*ij*_), we consider three different scenarios:

1. Absence of competition-induced death, where *p*_00_ = *p*_01_ = *p*_10_ = *p*_11_ = 0.
2. Presence of heterogeneous competition-induced death, where *p*_01_ and *p*_10_ are randomly sampled while meeting *p*_01_ + *p*_10_ + *X <* 1.
3. Presence of homogeneous competition-induced death, where *p*_00_ = *p*_01_ = *p*_10_ = *p*_11_*≠* 0.

When we randomly sample the huge parameter space (Table 2) by generating up to 10^6^ random parameter configurations, we found that only around 20.3% cases reach non-trivial stationary regimes in each of the above three direct competition scenarios (Figure 1). Therefore, direct competition does not make a huge influence on the probability for the bacterial population to persist (with or without the plasmid). The probability of coexistence increases under the absence of direct competition, while bistationary states only appear under the strain-dependent direct competition (0.15% of all cases).

**Figure 1:**
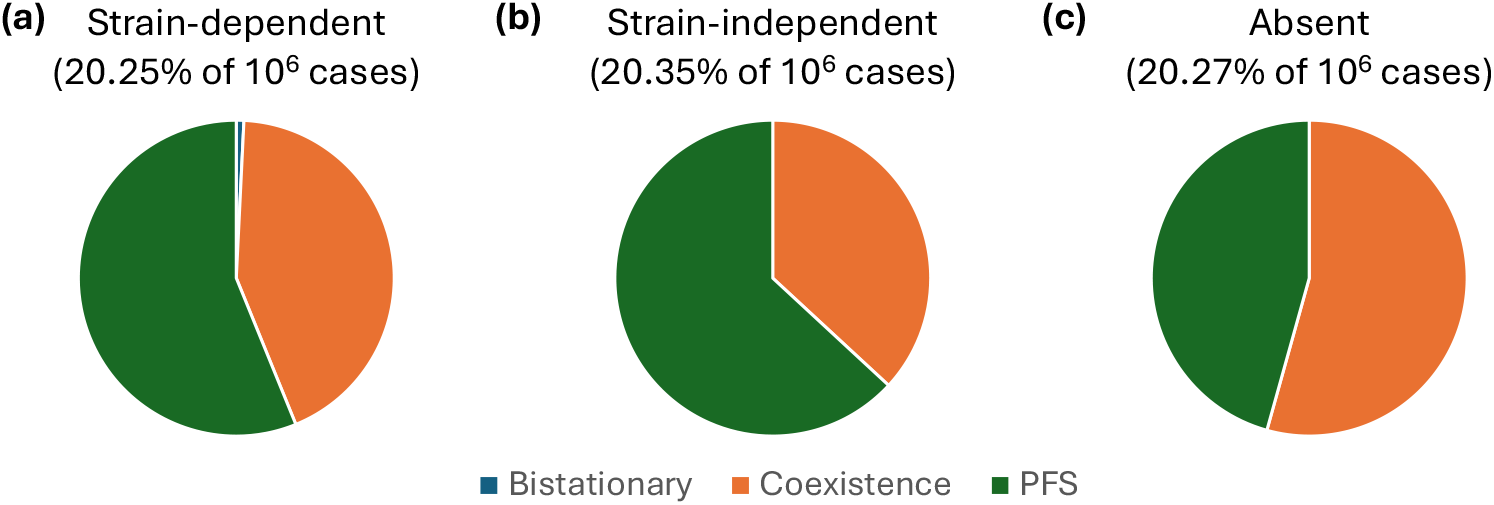
Non-trivial stationary regimes of a 1*B*1*P* system from 10^6^ iterations, where the direct competition is either (a) strain-dependent, (b) strain-independent, or (c) absent.

Starting from the reference parameter configurations where the direct competition is strain-specific vs. absent (Table 3), we first explore the dynamic regimes across the *β*_0_-*δ*_0_ (bacterial growth-death) plane (Figure 2). Low *δ*_0_ and moderate-to-high *β*_0_ favor coexistence and bistationary states under strain-specific direct competition (Figure 2a), while these parameter configurations only result in coexistence under the absence of direct competition (Figure 2b). As *δ*_0_ increases, the system tends toward plasmid-free states then extinction. When the direct competition results in bistationary state (red cross marked in Figure 2a, Table 3), the stationary state depends on the initial state: a low initial abundance of plasmid-carrying strain (*x*_1_) and a moderate-to-high abundance of plasmid-free strain (*x*_0_) favor plasmid-free state, while the rest combinations favor coexistence dominated by plasmid-carrying strain (Figure 3a-b). Notably, while the long-term numerical simulation only reflects stable stationary points (i.e. basin of attraction), the zero-growth isocline also reflects unstable, non-trivial stationary points (Figure 3c), where the strain dynamics might be temporarily attracted to before eventually attracted to the nearest stable stationary points.

**Table 3:** Model parameter values used in Figure 2a (the 3rd column) and Figure 2b (the 4th column). Note that direct competition is either strain-specific or absent, and that all the parameters reported here are the ones before being scaled by *γ*_0_ = 3.0.

|  |  |  |  |
| --- | --- | --- | --- |
| Strain-specific death rate | $\delta_0$ | (0, 10.0) | (0, 10.0) |
| Stress-induced death rate | $\delta_1$ | 1.0 | 1.0 |
| Cell division rate | $\beta_0$ | (0, 10.0) | (0, 10.0) |
| Segregation error probability | $\epsilon$ | 0.006 | 0.006 |
| Plasmid transmission probability | $\chi$ | 0.06 | 0.06 |
| Direct competition | $\begin{bmatrix} p_{00} & p_{01} \\ p_{10} & p_{11} \end{bmatrix}$ | $\begin{bmatrix} 0.44 & 0.37 \\ 0.56 & 0.28 \end{bmatrix}$ | $\begin{bmatrix} 0 & 0 \\ 0 & 0 \end{bmatrix}$ |
| Plasmid reproduction cost | $c$ | 0.2 | 0.2 |
| Plasmid resistance | $r$ | 0.6 | 0.6 |

**Figure 2:**
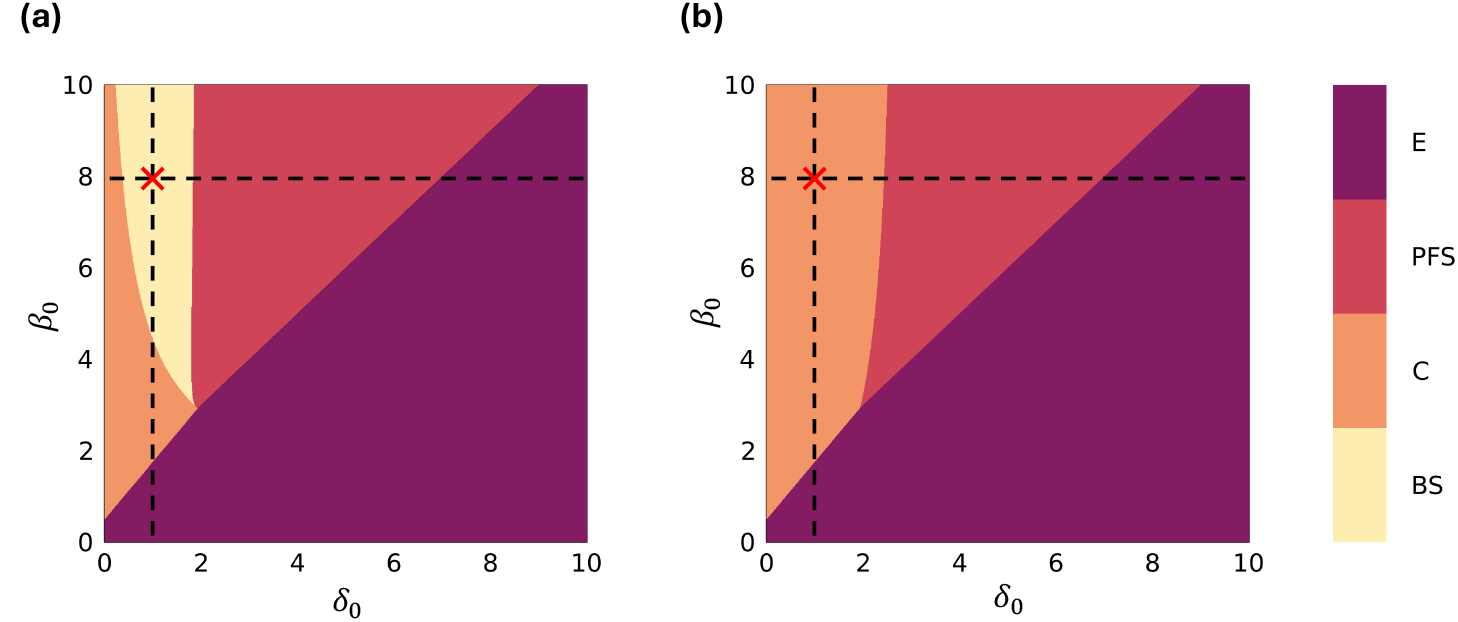
Stationary regimes of a 1*B*1*P* system across *β*_0_-*δ*_0_ plane where the direct competition is (a) strain-specific vs. (b) absent. E = extinction, PFS = plasmid-free state, C = coexistence, BS = bistationary state. The red crosses mark the reference parameter configurations (Table 3). The basin of attraction of the red cross in (a) is displayed in Figure 3.

**Figure 3:**
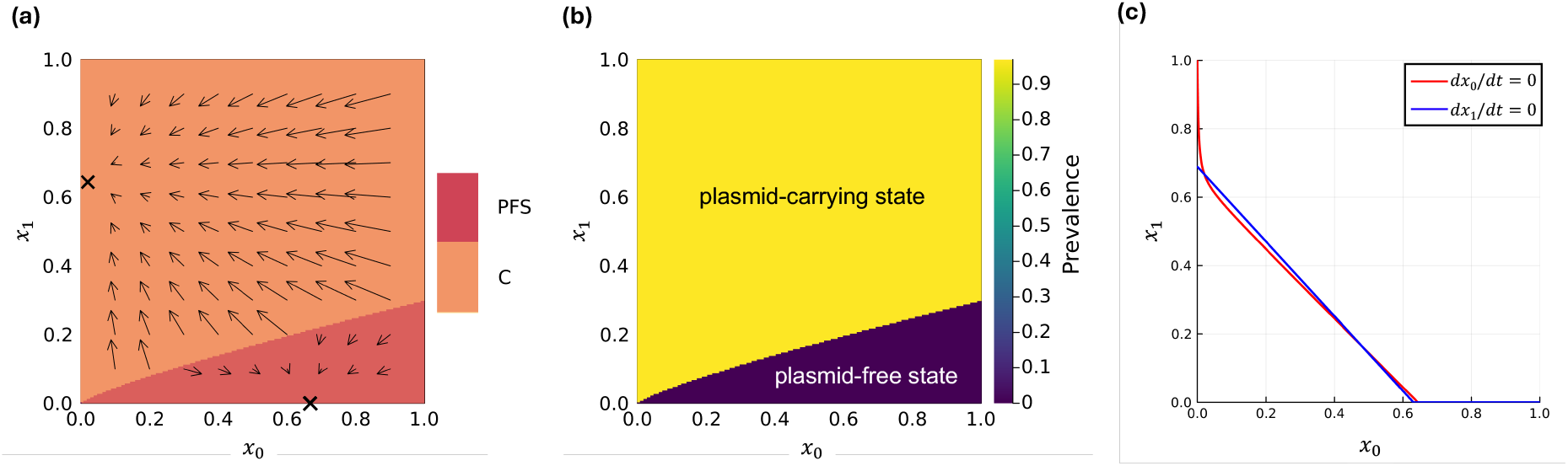
(a) Stationary regimes (E = extinction, PFS = plasmid-free state, C = coexistence), (b) plasmid prevalence after a very long time, and (c) zero-growth isocline across *x*_0_-*x*_1_ plane of a 1*B*1*P* system, based on the reference parameter configuration (red cross in Figure 2a, Table 3) and numerical simulation (*t* = 50000, *K* = 10^6^, *γ*_0_ = 3.0). The arrows and crosses in (a) denote the trajectories of population dynamics and the basin of attraction (i.e. stable stationary points). The three intersections in (c) are non-trivial stationary points (along the *x*_0_ axis): stable coexistence dominated by *x*_1_, unstable coexistence dominated by *x*_0_, and stable plasmid-free state. PFS = plasmid-free state, C = coexistence.

### Plasmid eradication across competition scenarios

Based on the reference parameter configurations (red crosses in Figure 2a-b, Table 3), we generate bifurcation diagrams and examine fast vs. slow hysteresis cycles of strain dynamics (initial abundances *x*_0_ = *x*_1_ = 0.01, corresponding to plasmid prevalence of 0.5; *K* = 10^6^, *γ*_0_ = 3.0) across ranges of a control parameter (*β*_0_, *δ*_0_, and *χ*) where direct competition is either strain-specific or absent (Figure 4, Figure 5). These analyses aim to assess whether the existing plasmid-carrying strain (*x*_1_) can be eradicated through targeted intervention strategies: temporally (1) decreasing the population growth rate *β*_0_, (2) increasing the death rate *δ*_0_, or (3) decreasing the conjugation rate *χ*, each applied at either a fast or slow pace (Table 4).

**Table 4:**
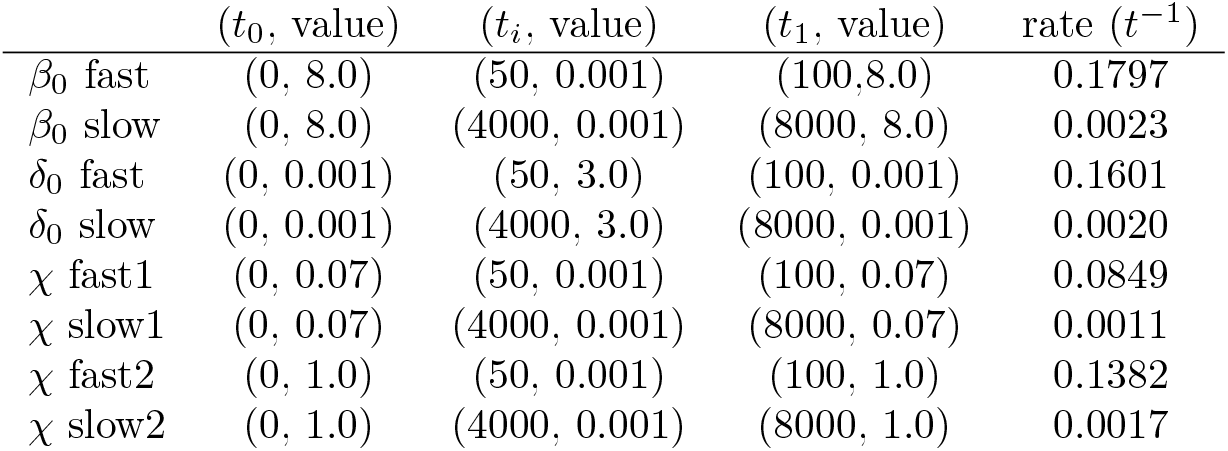
Time, parameter values, and rate of change used in the hysteresis cycles in Figure 4 and Figure 5. Note that *X* fast1 and *X* slow1 are used under strain-specific direct competition (Figure 4), while *X* fast2 and *X* slow2 are used under the absence of direct competition (Figure 5). Both the time and the parameters reported here are the ones before being scaled by *γ*_0_ = 3.0.

**Figure 4:**
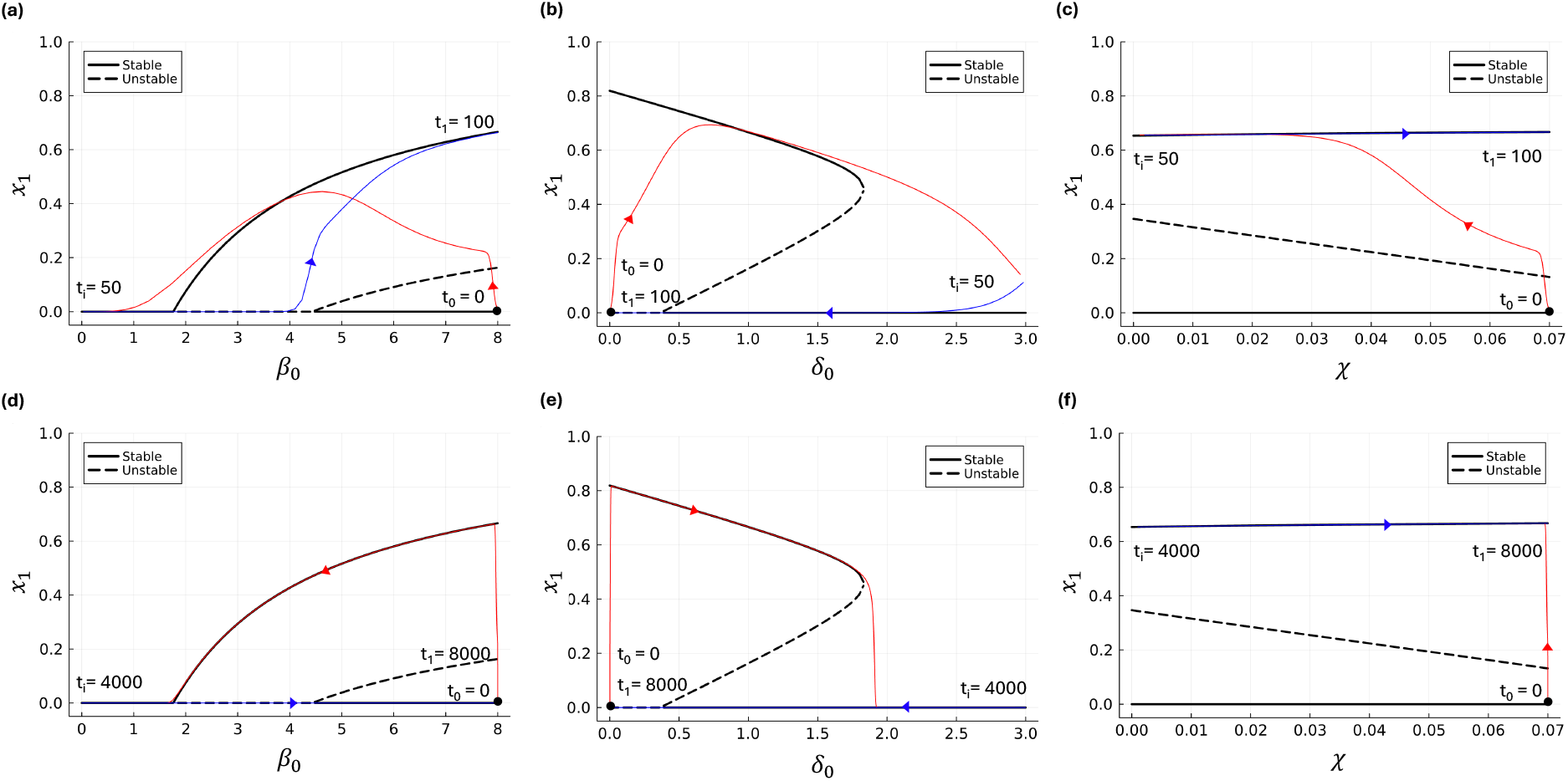
Bifurcation diagram and hysteresis cycles in terms of control parameters *β*_0_ (a, d), *δ*_0_ (b, e), and *X* (c, f) showing plasmid-carrying strain dynamics under strain-specific direct competition. Response to a fast (a-c) vs. slowly (d-f) decreasing/increasing parameters from high/low to low/high values (red line) and back (blue line). The black solid dots mark the starting points of the cycles.

**Figure 5:**
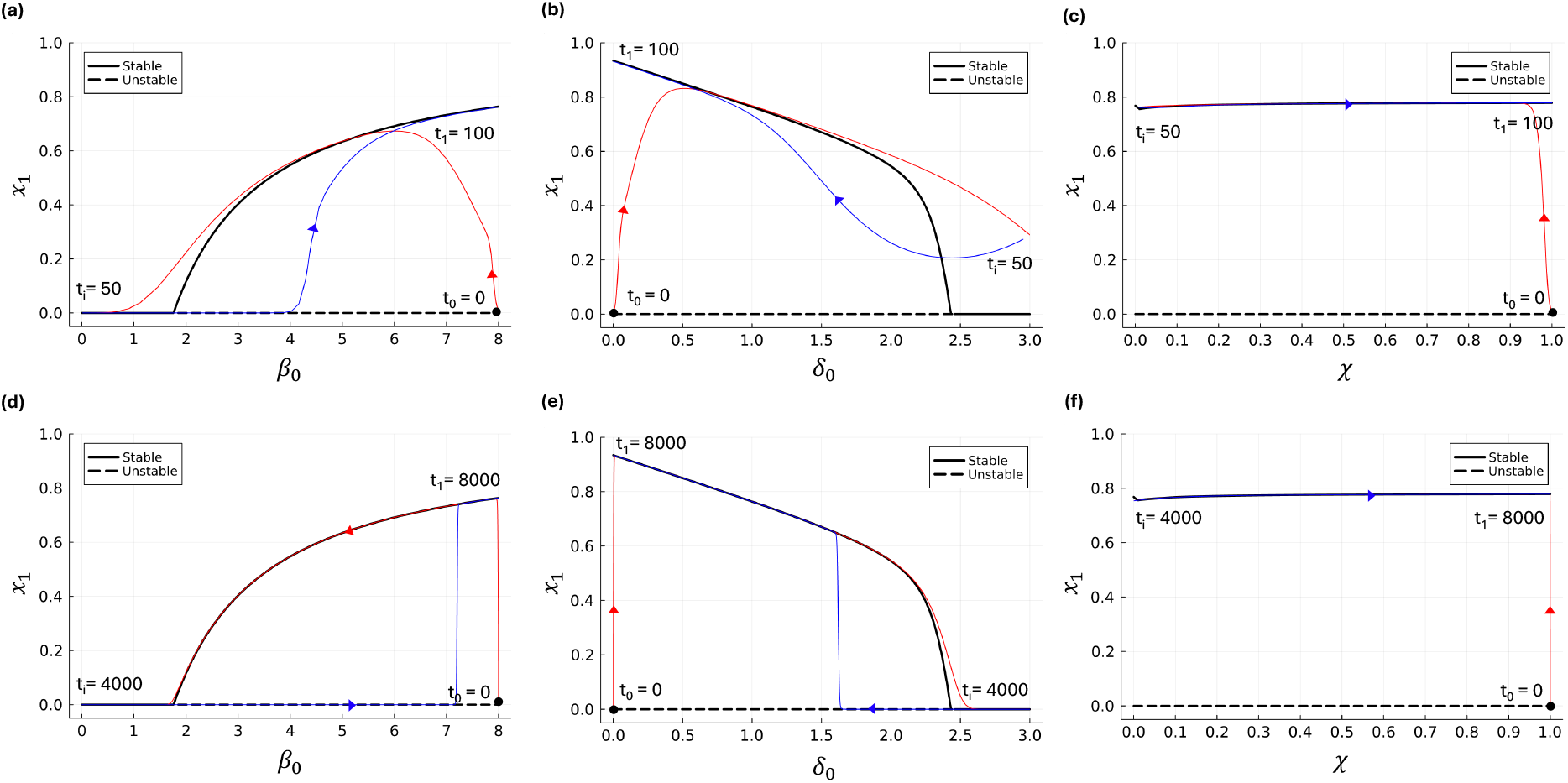
Bifurcation diagram and hysteresis cycles in terms of control parameters *β*_0_ (a, d), *δ*_0_ (b, e), and *X*(c, f) showing plasmid-carrying strain dynamics in the absence of direct competition. Response to a fast (a-c) vs. slowly (d-f) decreasing/increasing parameters from high/low to low/high values (red line) and back (blue line). The black solid dots mark the starting points of the cycles.

Under strain-specific direct competition, increasing *δ*_0_ is the most effective strategy: both fast and slow interventions successfully prevent the rebound of the plasmid-carrying strain during recovery of *δ*_0_ (Figure 4b, e). Decreasing *β*_0_ is only effective when applied slowly (Figure 4d); at a fast pace, the plasmid-carrying strain rebounds as *β*_0_ reaches half-recovery (Figure 4a). Decreasing *χ* fails to eliminate the plasmid-carrying strain at either pace, with the strain quickly reaching a stably dominant state even before *χ* begins to recover (Figure 4c, f).

In the absence of direct competition, all strategies fail to eradicate the plasmid-carrying strain (Figure 5a-f). A slow decrease in *β*_0_ (in comparison with a fast decrease) makes the rebound during recovery occur at larger values of *β*_0_, but does not prevent the strain from becoming stably dominant (Figure 5a, d). Similarly, a slow increase in *δ*_0_ (in comparison with a fast increase) makes the rebound during recovery occur at larger values of *δ*_0_ (Figure 5b, e); however, the plasmid-carrying strain becomes stably dominant before *δ*_0_ is even halfway recovered (Figure 5b).

In summary, the presence of strain-specific direct competition enables bistability and is key to successful plasmid eradication via interventions.

## 4 Discussion

In this study, we developed and analyzed a minimal one-bacterial population-one-plasmid (1B1P) model to investigate how the structure of direct competition between plasmid-free and plasmid-carrying strains shapes population dynamics. Our key finding is that the competition structure alone can fundamentally alter the systems behavior governing whether plasmid-carrying strains can invade, persist, or be eradicated. We identified three classes of equilibrium and derived the conditions for plasmid invasibility and eradicability, demonstrating that direct competition plays a pivotal role in both preventing plasmid invasion and enabling its eradication.

Notably, in the absence of direct competition, the system exhibits smooth and continuous transitions between states, enabling broader conditions for plasmid invasion and eventual dominance. In contrast, introducing strain-specific direct competition narrows the invasion window and can generate bistability and hysteresis dynamics where small changes in parameters or initial conditions trigger abrupt regime shifts between plasmid persistence and extinction. These non-linear behaviors have important implications for intervention strategies: the ability to prevent the establishment of plasmid-encoded antibiotic resistance depends on both the conjugation rate and the underlying competition structure, while successful eradication may hinge on whether the system resides in a bistationary regime, which is possible only under strain-specific direct competition. A key and novel insight from our work is that the structure of pairwise competition alone—independent of other ecological or evolutionary factors—is sufficient to induce qualitatively distinct dynamic regimes. These nonlinear behaviors carry important implications for intervention strategies: the success of efforts to prevent or eliminate plasmid-encoded antibiotic resistance depends not only on the type or strength of the intervention, but also critically on the underlying ecological interactions that govern competition.

Our findings have broad implications for understanding plasmid dynamics in both ecological and clinical microbial communities, emphasizing the critical role of competition structure, particularly direct interference competition. In environments such as the gut, soil, or other microbial habitats, bacteria frequently engage in both indirect exploitative competition (e.g., resource depletion that slows down cell division) and direct interference (e.g., toxin production that induces cell death) [15, 16]. Although these competitions have not been compared empirically, simulations have suggested that direct interference competition amplifies population differences more quickly than indirect exploitative competition [23]. Our model indicates that strain-specific direct competition can shift the system from coexistence regimes to bistationary regimes, where long-term outcomes are highly sensitive to initial conditions and changing parameters. This insight suggests that intervention strategies that overlook the ecological context—especially the form of inter-strain competition—may fail to anticipate scenarios in which plasmid-carrying strains persist. For example, outside of a bistationary regime, even temporarily increasing the mortality of both plasmid-carrying and plasmid-free strains (e.g., via a broad-spectrum antibiotic to which neither is resistant) may be insufficient to eliminate the plasmid, as the system may remain trapped in a plasmid-carrying state. The predicted pattern of regime shifts and hysteresis is therefore of particular concern: in the absence of hysteresis, plasmid-carrying strains can persist or re-emerge when selective pressures are removed, raising the risk of relapse or reestablishment of antibiotic resistance. In empirical trials, patterns of hysteresis may serve as indirect indicators of underlying direct competition structure, potentially providing early warnings of approaching ecological and clinical tipping points.

The persistence of plasmids in microbial populations has traditionally been explained through a combination of mechanisms such as high transmission rates, host-specific plasmid traits, positive epistasis, source-sink transmission mode, compensatory evolution, and adaptive evolution [24]. However, many models in the literature assume either well-mixed populations or homogeneous interactions between hosts, and often neglect the competition structure that shapes community dynamics. When competition is considered, it is typically modeled implicitly through resource- or density-dependent growth, without accounting for its structure or strain specificity (e.g. [11, 12]). In contrast, our 1B1P model demonstrates that the structure of direct, strain-specific competition alone can determine whether plasmids invade, persist, or are eradicated even in the absence of differential fitness, selection, or stochasticity. This focus adds a new ecological dimension to understanding plasmid stability by revealing that bistability and hysteresis can emerge solely from how plasmid-carrying and plasmid-free strains interact competitively. Such nonlinear outcomes have not been predicted by traditional microbe-plasmid models, though they have been acknowledged in other models, such as those describing microbial cross-feeding and vector-borne disease transmission [25, 26]. By embedding plasmid transmission within an explicit direct competition network, our work complements existing theories by showing that the topology and asymmetry of inter-strain interference can profoundly affect long-term system outcomes. Our findings underscore that ecological interaction structure may act as a critical constraint or facilitator of antibiotic resistance control. Integrating this ecological perspective with existing models will be crucial to developing a better understanding of plasmid ecology and designing robust intervention strategies.

Our minimal 1B1P model offers essential insights into the influence of direct competition on plasmid dynamics, but overlooks several biological complexities. It presumes a uniform environment, neglecting spatial structure, environmental fluctuations, and host immune responses, all of which play critical roles in bacterial interactions. Notably, spatial structure is crucial for improving predictions and management strategies concerning the dissemination of plasmid-mediated antibiotic resistance genes among human populations. In this context, our model can be used as the building block of a metapopulation model (e.g. [27]), where each human individual hosts a growing bacterial infection that can transmit to others, posing a risk of spreading antibiotic resistance. Moreover, our model does not account for the diversity of plasmids and bacteria, nor the possibility of co-infection by multiple plasmids. It also excludes evolutionary and stochastic processes, despite the fact that interference strength and other bacterial and plasmid traits can evolve, and that demographic stochasticity can arise. These factors may lead to divergent outcomes across ecological contexts (e.g., [28, 29]). Future research endeavors should aim to extend this model to incorporate bacterial and plasmid diversity, spatial structure, stochasticity, and evolutionary dynamics impacting both host and plasmid traits. Validation efforts, using synthetic communities or actual biological systems, will be essential to verify model predictions and improve intervention strategies in realistic ecological settings.

In summary, our study reveals that the structure of direct, strain-specific competition alone can drive qualitatively distinct plasmid dynamics, including bistability and hysteresis, with critical implications for the persistence and control of plasmid-mediated antibiotic resistance. Small changes in interaction parameters can drive disproportionate effects—echoing the broader theory of tipping points in ecology. By demonstrating that such nonlinear outcomes emerge independently of other ecological or evolutionary mechanisms, our work highlights competition structure as a key and underappreciated determinant of intervention success. These findings emphasize the need to integrate ecological interaction networks into models of plasmid transmission and antibiotic resistance. Future efforts to design effective interventions must account not only for selective pressures but also for the competition structure that shapes microbial community dynamics.

